# Connections of early visual areas with posterior parietal and temporal cortex in galagos, a strepsirrhine primate

**DOI:** 10.64898/2025.12.27.696608

**Authors:** Qimeng Wang, Jon H. Kaas, Iwona Stepniewska

## Abstract

To better understand the cortical connections and organization of visual areas in galagos, we examined the interconnections among cortical visual fields and their relationships with posterior parietal cortex (PPC) and temporal regions associated with dorsal and ventral streams of visual processing. In five galagos, two to four distinguishable tracers were injected into different visual areas, allowing direct comparison of connection patterns within the same cases. To reveal distributions of labeled neurons for each injection, labeled cells were plotted from serial brain sections cut parallel to the flattened cortical surface and summed across sections to generate surface reconstructions. Alternate sections processed for cytoarchitectonic features were used to identify cortical borders, especially those of V1 and middle temporal visual area (MT). Overall, our results support the conclusion that regions of V2 and V3 represent the contralateral visual hemifield in parallel with V1 and with each other. However, dorsal V3, representing the lower visual hemifield, includes at least one discontinuity where representations of the upper visual field extend to the dorsal border of V2. This portion of V3 appears to belong to the dorsomedial visual area (DM), which extends rostrally from V2 into PPC. The dorsal part of the DL-V4 region receives projections from other parts of dorsolateral visual area (DL), central V1, V2 and V3, inferotemporal (IT) cortex, the MT complex, and PPC regions surrounding the intraparietal sulcus (IPS). More central portions of DL-V4 receive inputs from central representations of V1, V2, and V3, as well as from PPC regions lateral to the IPS, the MT complex, and upper IT cortex. The ventral part of DL receives projections from central V2, caudal PPC adjoining DM and ventral PPC, and from IT cortex. These patterns indicate that the DL-V4 region serves as a major node linking dorsal and ventral streams and likely includes more than one functionally distinct visual area. In addition, areas MT and DM show strong reciprocal connections with PPC, while the connections of IT cortex indicate that much of this region is visual in nature having strong connections with higher order visual areas and it is composed of multiple functionally specialized visual domains.

**Key points:**

1. Organization of the visual cortex in galagos is much like that in New World and Old Word monkeys.
2. Patterns of cortical connections of early visual areas V1 and V2 support the view that dorsal V3 has a gap in the representation of the lower visual field that is occupied by the proposed dorsomedial visual area (DM).
3. Visual areas DM and middle temporal visual area (MT) provide the major visual inputs to posterior parietal cortex (PPC) of the dorsal stream of visual processing for actions.

## Introduction

In the present study, we injected two to four distinguishable tracers into visual cortical areas or regions of five galagos, a strepsirrhine primate often considered to resemble early primate ancestors. Our aim was to better understand the areal organization of early stages of cortical visual processing in galagos and to determine how these areas contribute to the proposed dorsal and ventral visual processing streams that extend into posterior parietal cortex (PPC) and inferior temporal cortex (IT).

Early euprimates emerged around the Cretaceous–Paleogene boundary, approximately 66 million years ago, and subsequently diversified into the major branches of the primate radiation that include the extant primate lineages today (Steiper and Seiffert, 2012; Silcox and Lopez-Torres, 2017). A fundamental question in primate evolution concerns how different taxa resemble one another and how they differ in terms of cortical organization and specialization. Although brain size and neocortical complexity vary greatly across primate species (Dunbar and Shultz, 2007; Powell et al., 2017), the basic plan of cortical sensory and motor fields appears conserved. Among extant strepsirrhines, galagos and mouse lemurs are thought to most closely resemble early euprimates, which were small, nocturnal mammals highly adapted for visually guided locomotion among the fine branches of tropical forests (Kaas et al., 2022).

Despite their relatively small brains, galagos possess most of the visual cortical areas identified in New World and Old World monkeys, apes, and humans. These include the primary visual area (V1), the second visual area (V2), and the middle temporal area (MT), all of which are consistently identified across primate species. However, the definition and boundaries of other proposed visual areas remain controversial. The third visual area (V3) was originally proposed as a mirror-image representation of V2, based on early studies of cortical connections (Cragg, 1969; Zeki, 1969; Gattass et al., 1988). Yet subsequent research raised doubts about the continuity of dorsal V3 (V3d), which represents the lower visual field, and its relationship to ventral V3 (V3v), which represents the upper field. Some investigators proposed that V3d be replaced by several smaller visual areas, including the dorsomedial (DM) and medial (M) visual areas (Allman and Kaas, 1975; 1976). Others have proposed modified V3d configurations incorporating interleaved representations of adjacent fields such as M (Rosa et al., 2005; Buckner and Yeo, 2014). Similarly, a dorsolateral visual area (DL) was proposed to occupy the region between central V2 and MT (Allman and Kaas, 1974), while other studies have referred to this region as V4 (Zeki, 1971). The dorsal and ventral boundaries of DL–V4 remain uncertain, and the area may include rostral (DLr) and caudal (DLc) subdivisions (Cusick and Kaas, 1988). Alternative schemes have extended V3 and V4 along the ventral border of V2, with shorter extensions along its dorsal border (Kaas, 1997; Kaas and Lyon, 2001; Lyon and Kaas, 2002a; Lyon and Connolly, 2011; Rosa et al., 2005; Stepniewska et al., 2005). These variations in proposed cortical maps underscore the challenge of defining a consistent model of visual cortex organization across primates and highlight the need for comparative studies in strepsirrhine species. One goal of the present investigation, therefore, was to clarify the organization of visual cortical fields in galagos through anatomical tracing of intrinsic and interareal connections.

A second goal was to better define how early visual areas contribute to the dorsal and ventral streams of higher visual processing. Early lesion and behavioral studies in macaques established the concept of two major cortical processing streams: a ventral “what” pathway for object identification and a dorsal “where” pathway for spatial and visuomotor processing (Ungerleider and Mishkin, 1982). Later refinements emphasized the role of the dorsal stream as a “how” pathway, mediating visually guided actions such as reaching and grasping (Goodale and Milner, 1992). While these pathways have been extensively characterized in macaques, their organization in New World monkeys and strepsirrhines remains less well understood. Nevertheless, domains within posterior parietal and motor cortices that mediate distinct action types appear remarkably conserved across galagos, squirrel monkeys, and macaques (Kaas and Stepniewska, 2016). The primary visual inputs to these two streams are thought to originate from area MT for the dorsal stream and from the DL-V4 complex for the ventral stream (Felleman and Van Essen, 1991; Kaas et al., 2022).

By analyzing the results of multiple tracer injections in the visual cortex of galagos, the present study identifies the sources and patterns of cortical connections with the caudal half of PPC and IT. These findings provide new insights into both the areal organization of visual cortex and the early evolutionary foundations of the dorsal and ventral visual processing streams in primates.

## Materials and Methods

Five adult galagos (*Otolemur garnetti*) were used in this study. Anatomical tracers were placed in the presumptive DL and adjacent regions to reveal cortical connections. All experimental procedures followed the Guide for the Care and Use of Laboratory Animals established by the National Institutes of Health and were approved by the Animal Care and Use Committee of Vanderbilt University.

### Surgery and tracer injection

Galagos were initially anesthetized with 2% isoflurane inhalant anesthetic or an intramuscular injection of ketamine hydrochloride (30 mg/kg) and were maintained a surgical level of anesthesia during the procedure. Surgical procedures were carried out under aseptic conditions with the animal’s vital signs being consistently monitored. After animals were fixed on the stereotaxic apparatus, a craniotomy and durotomy were performed on one hemisphere to expose lateral occipital, temporal and posterior parietal cortices in the approximate DL region, posterior to the caudal tip of IPS, and upper temporal visual cortex. The cortical surface was photographed and printed to allow documentation of injection sites of tracers.

In each galago, two to four different anatomical tracers, including Fluororuby, Fluoroemerald (10% FR or FE in distilled water), Diamidino Yellow (2% DY in phosphate buffer), Fast Blue (3% FB in distilled water) and Cholera Toxin subunit B (1% CTB in distilled water) were placed at different locations in DL-V4 region and adjacent visual cortex for a total of 16 injections. Pressure injections of tracers were made with Hamilton syringes at two depths, 1.6 mm and 1 mm, below the pial surface to include the whole cortical thickness. After injections, the exposed cortex was covered with gelfilm, and the opening in the cranium was covered with a thin piece of hard dental cement fixed to the surrounding skull before the skin was sutured over the opening. Animals were given injections of antibiotics and placed in a recovery cage under careful monitoring until fully awake. After recovery, they were returned to home cages.

### Tissue processing and histology

Five to nine days after surgery, galagos were administered a lethal dose of sodium pentobarbital (120 mg/kg), and then perfused intracardially with 0.9% phosphate-buffered saline (PBS), followed by 2% paraformaldehyde in phosphate buffer (fixative) and 10% sucrose in fixative. The brain was removed from the skull and bisected into two hemispheres. Left and right cortices were then separated from subcortical structures and underlying white matter, and manually flattened to unfold sulci before squeezed between two glass slides. After two hours of post fixation in the same fixative, the flattened cortices were transferred into 30% sucrose solution for cryoprotection and stored at 4°C overnight for microtomy.

Cortices were sectioned in parallel to the pial surface at a thickness of 40 μm and saved in four or five series. One series of sections was mounted without further processing for fluorescence microscopy. In cases in which CTB was used, one series of sections was processed to reveal CTB labeled cells (Angelucci et al., 1996). In all cases, one series of sections was processed for cytochrome oxidase (CO; Wong-Riley, 1979) and a second series for myelinated fibers (Gallyas, 1979) in order to reveal cortical architectures, particularly the well recognizable primary visual (V1) and middle temporal visual areas (MT).

### Data analysis and anatomical reconstruction

Neurons labeled with fluorescent substances were plotted using Igor Pro 3.14 (WaveMetrics Inc.) coupled with a Microcode II digital readout (Boeckeler Instruments Inc.) and a Leitz Orthoplan 2 microscope (Leica Microsystems Inc.) under 16× magnification under fluorescence illumination. Stereo Investigator 2019 (MBF Bioscience) coupled with a Zeiss Axio Imager 2 microscope (Carl Zeiss AG) was used to plot CTB-labeled neurons under 20× magnification, and to image CO (Figure 1a) and myelin sections under 5× magnification. For each case, three to four sections of each series were selected to cover across the superficial, middle and deep cortical layers. Plotted data with section outlines were imported into Adobe Illustrator CC2014 (Adobe Inc.) to be compiled into a surface-view reconstruction of distributions of labeled neurons, which was then aligned with images of architecture staining sections by matching common blood vessels and other anatomical landmarks to have information of area boundaries integrated. Identification of architectonic borders of cortical regions followed those of previous studies (Preuss and Goldman-Rakic, 1991; Wu and Kaas, 2003; Wong and Kaas, 2010). In cases where these areas were architechtonically indistinct, areal borders were estimated based on measurements from other cases and outlined with dashed lines.

**Figure 1.**
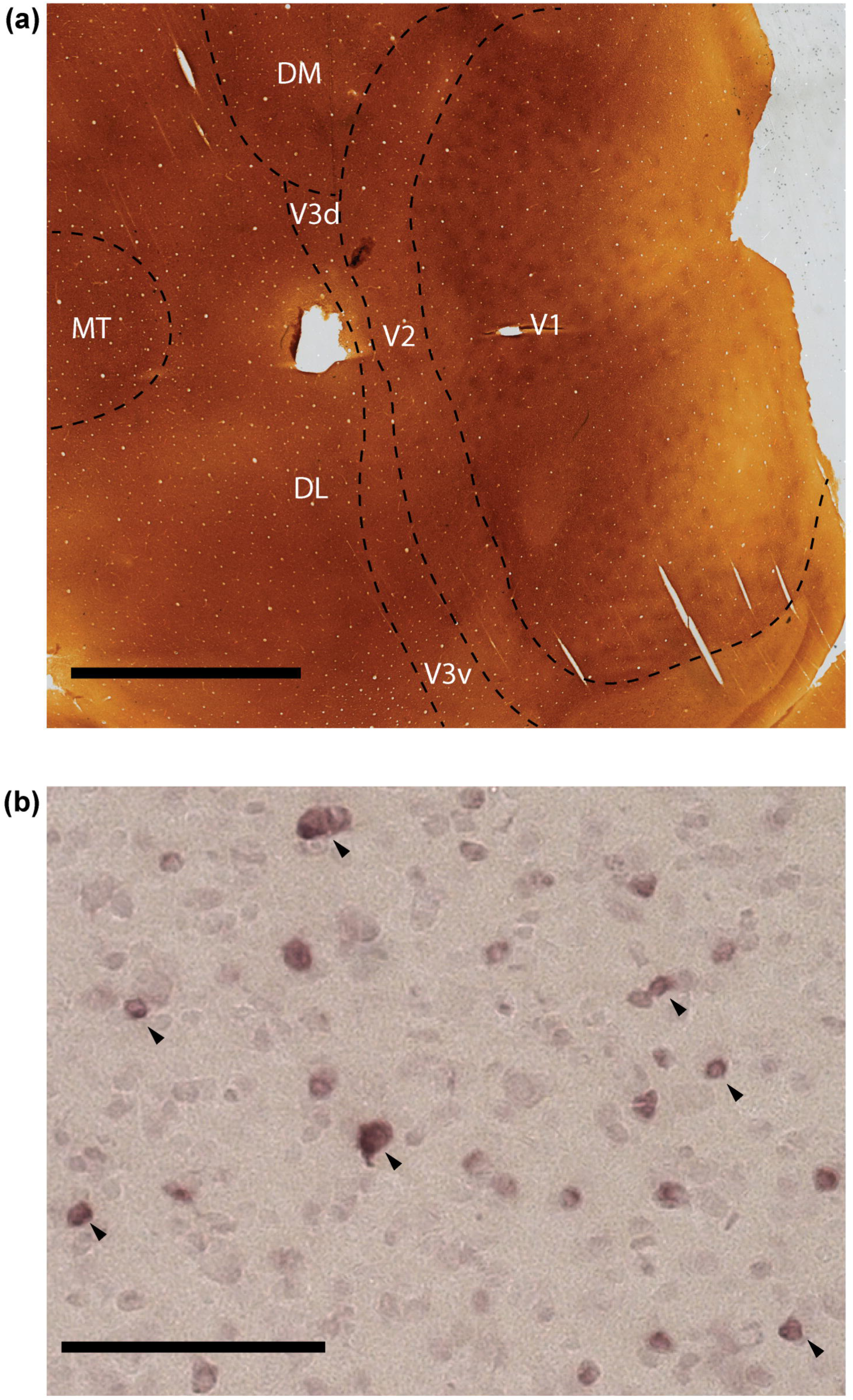
Photomicrographs of sections through the flattened visual cortex. (a) Section stained for CO with marked visual areas. V1, primary visual area; V2, second visual area; V3, third visual area (d, dorsal and v, ventral); DL, dorsolateral visual area; DM, dorsomedial visual area; MT, middle temporal visual area. Scale bar = 5 mm. (b) CTB labeled neurons (some indicated by arrows) in region of DL surrounding the injection site in case 03-40. Scale bar = 1 mm.

The flattened cortex preparation allows a direct visualization of surface-view cortical patterns of labeled cells and architectural features. It also avoids distortions inherent in reconstructions based on sections cut in standard planes. The flatten cortex preparation is therefore superior for accurately localizing boundaries of cortical areas and labeled neurons from a series of sections.

## Results

Our results are based on five galagos where up to four distinguishable tracers were placed in various locations in cortical visual areas or regions to identify especially connections of early visual areas with locations in PPC, reflecting connections devoted to the dorsal stream of higher visual processing, and connections with caudal inferior temporal cortex (ITc) devoted to the ventral stream of higher visual processing. The results reveal the locations of retrogradely labeled neurons projecting to the injection sites as these are more reliably identified than labeled axon terminals, although most projections are commonly bidirectional (Rockland, 2022). The patterns of labeled neurons are shown on surface views of flattened cortical hemispheres, which combines labeled neurons across cortical layers, as previously described (Rockland, 2022). These results are related to the proposed arrangements and histological distinctions of visual cortex in galagos.

### Case 03-40

In this case, injections of different tracers were placed in dorsal DL, central DL and ventral V2 (Figure 2). As expected, the ventral V2 injection (FR) had a somewhat elongated core along the medial to lateral injection track which mainly labeled neurons in the adjoining ventral portions of V2 and V1 where the paracentral upper visual field is represented (Figures 2a, 2c). Small patches of neurons were labeled in central V2 and V1. Another major location of labeled neurons was dorsomedial to the putative dorsal border of DL, possibly in the representations of the upper field of DM, which extends to the V2 border. Labeled neurons in the region of ventrolateral the crescent areas of the MT complex (MTc) suggest this portion of MTc represents the paracentral upper visual field. A tiny injection of DY in central DL, representing paracentral vision of the upper visual field, received inputs mainly from the ventral portions of V1, V2 and V3 (Figures 2a, 2b). Additional scattered inputs were from ventral DL, DM, MT, MTc and posterior IT region adjacent to V3. The CTB injection near the postulated dorsomedial border of DL labeled neurons over much of DL (Figures 1b, 2a, 2d), with much denser distributions in central and dorsal DL around the injection core than in ventral DL. Densely packed labeled neurons were observed in dorsal V2, along dorsal and ventral banks of IPS in caudal PPC (PPCc), as well as in ventral MST. Labeled neurons were also distributed in central V1, dorsal and ventral V3, ventral DM, as well as MT and its adjacent areas MTc and FST. The results suggest that much of the proposed territory of DL-V4 represents paracentral vision, and the lower margin of DM has connections with PPC along the caudal half of IPS. Some parts of DL have connections with separate parts of the inferior temporal cortex (IT), including parts that have upper field V2 connections.

**Figure 2.**
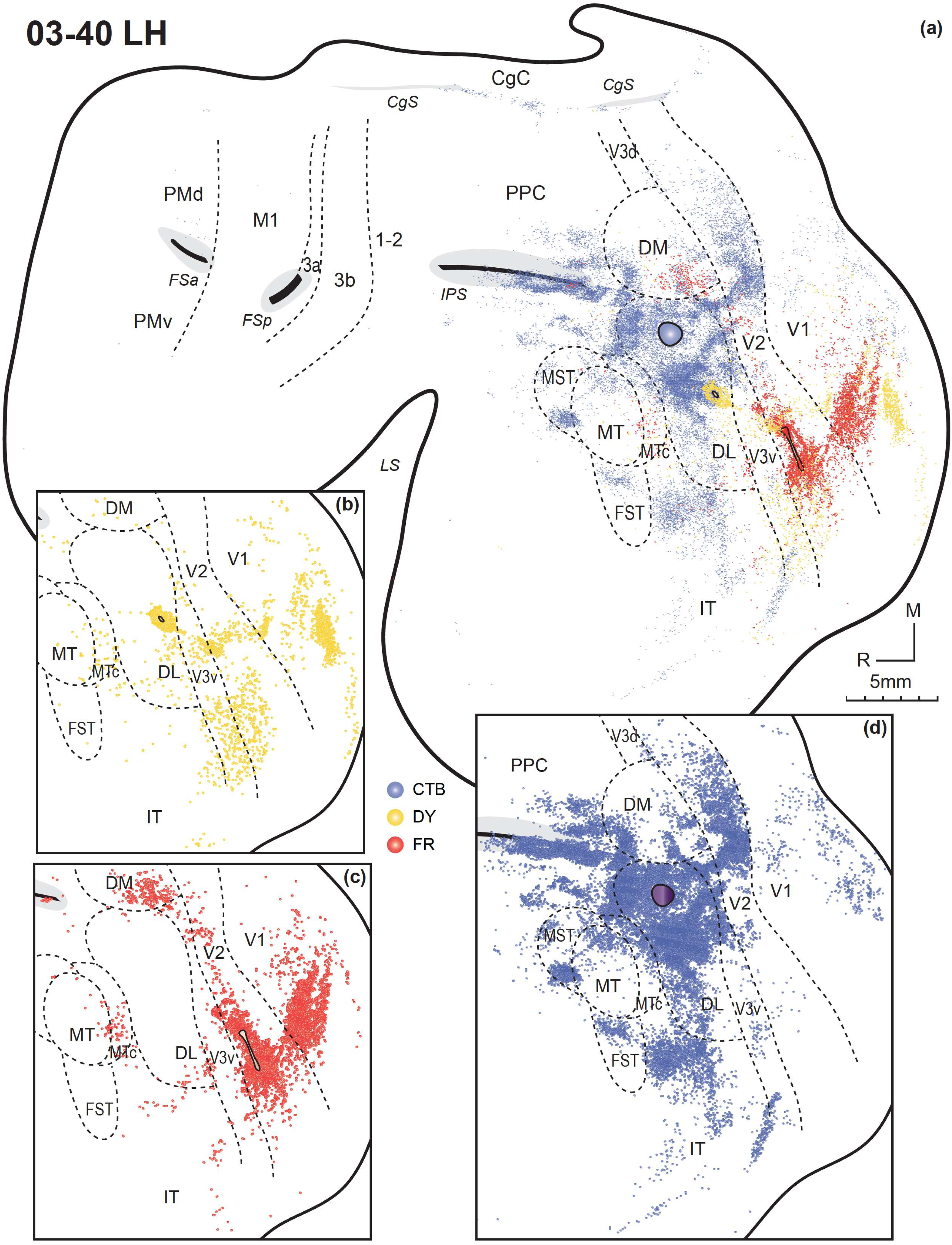
Distributions of labeled neurons after DY, FR and CTB injections in the left hemisphere of galago 03-40, shown altogether in the flattened cortex (a), as well as shown individually by tracers DY (b), FR (c), and CTB (d). Each dot represents a labeled neuron. The injection core of each anatomical tracer is outlined and filled with corresponding color. Areal boundaries are indicated by dotted lines. The approximate location of proposed visual areas, based on architectonic borders, most clear for V1 and MT, and estimated from previously proposed relationship, are shown as well as the expected location of somatosensory areas (3a, 3b and 1-2), primary motor cortex (M1) and dorsal (PMD) and ventral (PMV) premotor areas. Other references are for cingulate cortex (CgC), the cingulate sulcus (CgS) and the lateral sulcus (LS). Visual areas include V1, V2, dorsal and ventral V3, dorsolateral visual area DL, the MT, the MT crescent (MTc), the medial superior temporal area (MST), and the fundal area of the superior temporal sulcus (FST). The regions of posterior parietal cortex (PPC) and infratemporal cortex (IT) are indicated. M, medial; R, rostral. Scale bar = 5 mm.

### Case 03-18

This case received three injections in visual areas of cortex (Figure 3). A tiny injection of FE placed in dorsal V3 (V3d) near the proposed border with DL labeled neurons mainly in dorsal V1, representing the lower visual field (Figures 3a, 3c). Other labeled neurons were found in adjoining dorsal portion of V2 and V3. Labeled neurons were not detected in PPC or IT. A large injection of DY in the presumable location of central DL labeled central vision parts of both V2 and V1, as well as MT and FST regions (Figures 3a, 3d). Importantly, densely labeled cells were also detected in ventral portions of V2 and V3, in PPC just lateral to IPS, and some patches of labeled neurons were present in IT. A FR injection was along the injection track in IT which labeled neurons in surrounding regions of IT extending to FST, and in the ventral portion of PPC (Figures 3a, 3b). The results suggest that higher order visual regions of PPC and IT are directly interacting.

**Figure 3.**
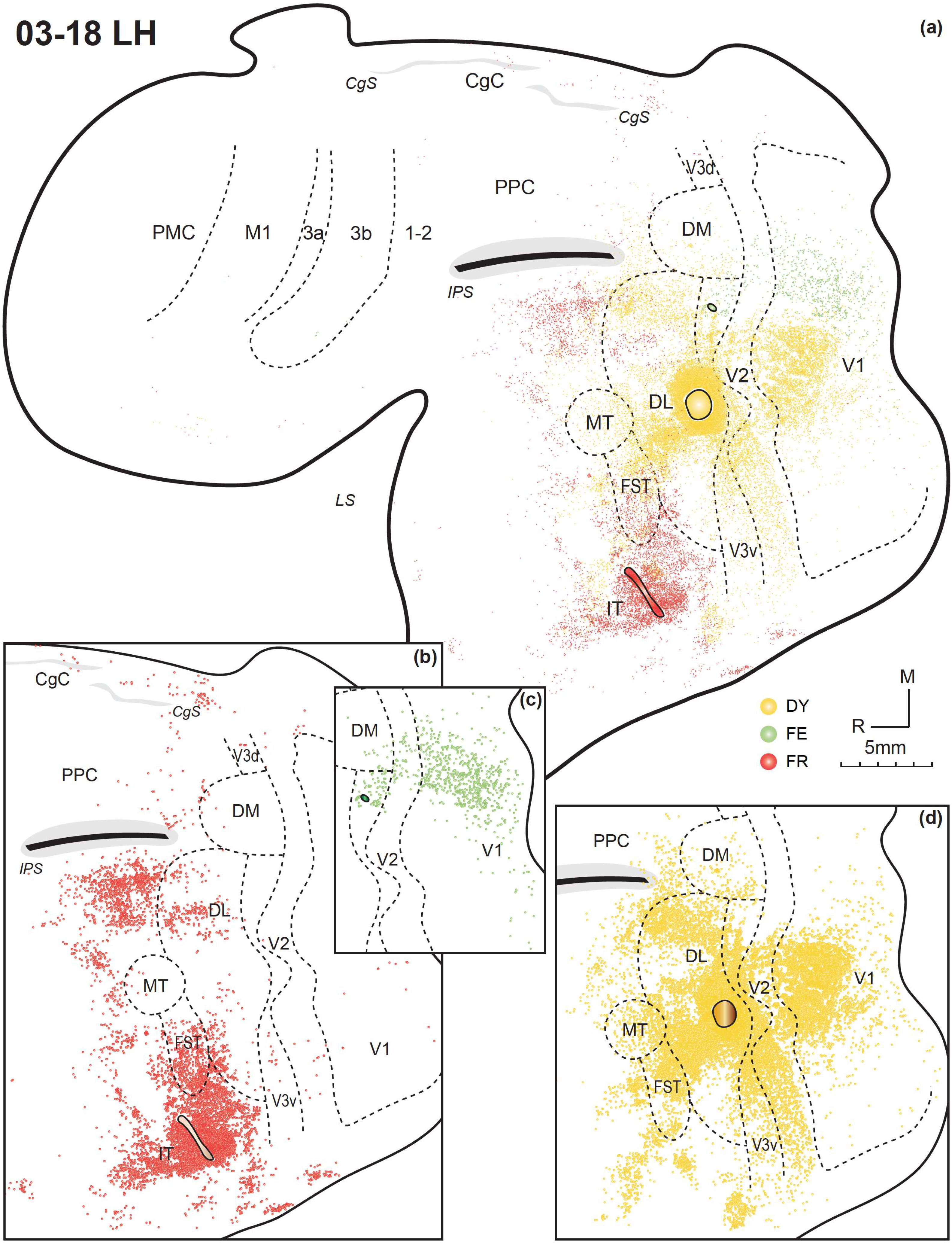
Distributions of labeled neurons after FR, FE and DY injections in the left hemisphere of galago 03-18, shown altogether in the flattened cortex (a), as well as shown individually by tracers FR (b), FE (c), and DY (d). Other conventions as in Figure 2.

### Case 00-78

In this case, two large (FB, DY) and two small (FR, FE) injections of four different tracers were placed dorsoventrally in higher order visual cortex (Figure 4). The FR injection was at the border of V2 and V3, representing paracentral vision with possible involvement of the upper visual field (Figures 4a, 4d). Besides densely labeled neurons in ventral V2, V3 and caudal parts of DL surrounding the injection site, another focus of label was present in the central part of V1. Some labeled neurons were distributed in other more medial parts of DL as well as in MTc and possibly in DM. The large FB injection site near the dorsomedial border of DL and presumably including ventral DM primarily labeled adjoining regions of cortex, including parts of PPCc, DM, DL and dorsal V2 (Figures 4a, 4b). Besides, parts of V2 and V1 that represent paracentral vision slightly more into the upper visual field contained dense patches of labels. Other labeled neurons were also observed in parts of MT, MTc and less in the middle superior temporal visual (MST), as well as locations in IT. The DY injection in the medial portion of DL labeled neurons in upper V2 and V1 (Figures 4a, 4e), suggesting the representation of the lower visual field. Other labeled patches were distributed in DL, caudal PPC, the MT complex and IT. The tiny FE injection site estimated in the region of ventral DL, labeled neurons in ventral V2, V3 and V1 that represent the upper visual field, and sparsely in the MT complex and IT (Figures 4a, 4c). The retinotopic patterns of connection revealed by the DY injection conform with the proposed retinotopy of DL, while connections revealed by the FB injection that are near the upper border of DL, are mainly from involvement of ventral DM.

**Figure 4.**
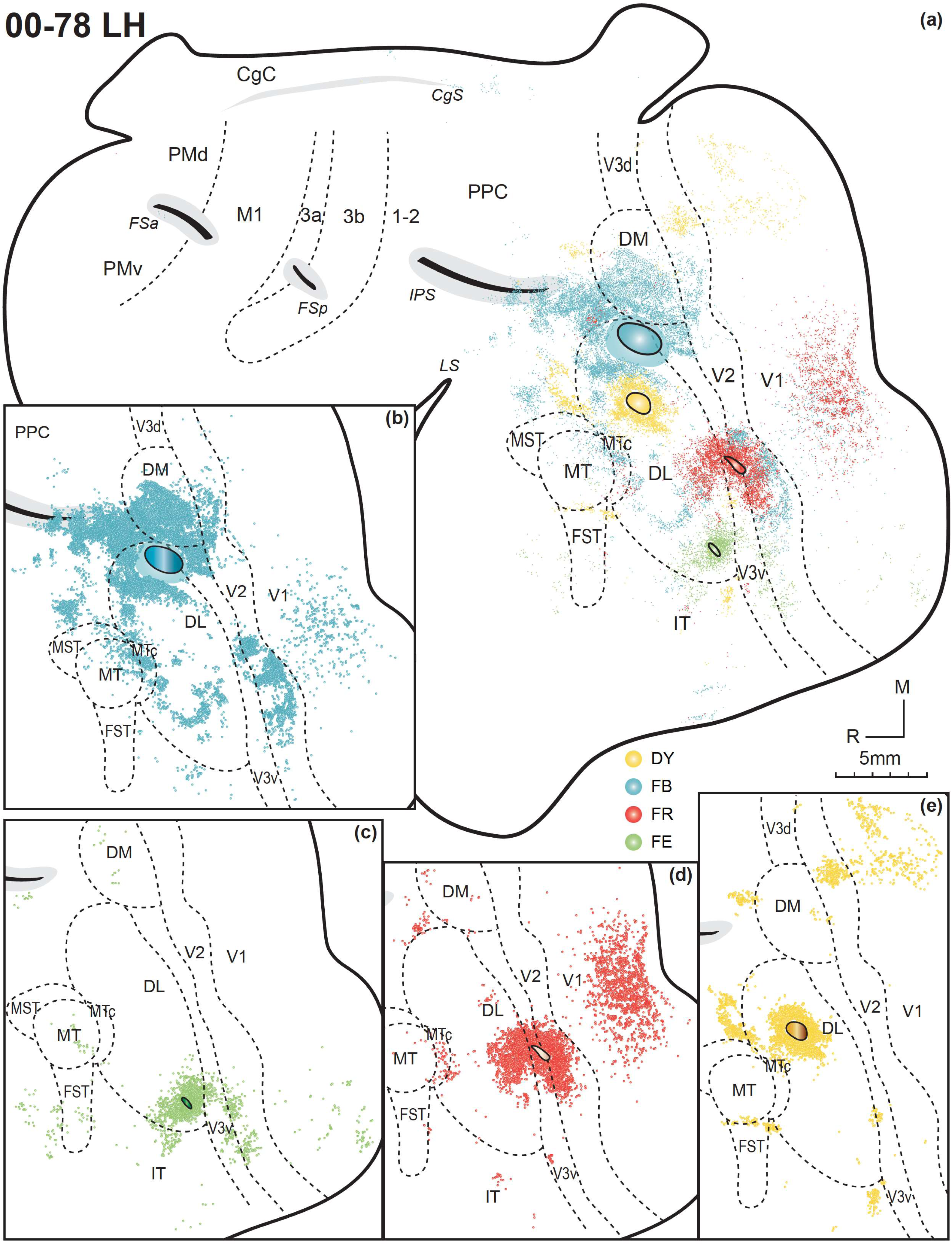
Distributions of labeled neurons after FB, FE, FR and DY injections in the left hemisphere of galago 00-78, shown altogether in the flattened cortex (a), as well as shown individually by tracers FB (b), FE (c), FR (d), and DY (e). Other conventions as in Figure 2.

### Case 00-55

Injections of four tracers were placed in visual-related areas of the left hemisphere in this case (Figure 5). FB that was injected in lateral MT representing the upper visual field labeled neurons in ventral V2 and V1 of paracentral vision of the upper field (Figures 5a, 5e). Major patches of labeled neurons were also seen in ventral V3 with encroachment on the ventrocaudal tip of DL, in the DM region next to dorsal V3, ventral PPC lateral to IPS, and the entire MT complex, including FST, MTc and MST. Thus, MT appears to receive inputs from other members of the MT complex, DM and PPC. CTB that was injected at the border of dorsal DL and MTc labeled surrounding neurons in dorsal DL and MTc (Figures 5a, 5d), with small clusters in MT, MST, FST and ventral DL. Some patches of neurons were labeled in dorsal V1 and V2 representing the lower visual field, while a group of neurons were labeled in the juncture of DL, ventral V2 and V3. The elongated FR injection at the ventral edge of DL labeled most of neurons in ventral DL, central V2, and PPC region ventrolateral to IPS (Figures 5a, 5b). A good amount of neurons ventral to DL in IT were also labeled, possibly because the injection was close to DL-IT border. A tiny FE injection placed in dorsal IT adjacent to ventral DL only labeled a few neurons close to the injection site (Figures 5a, 5c).

**Figure 5.**
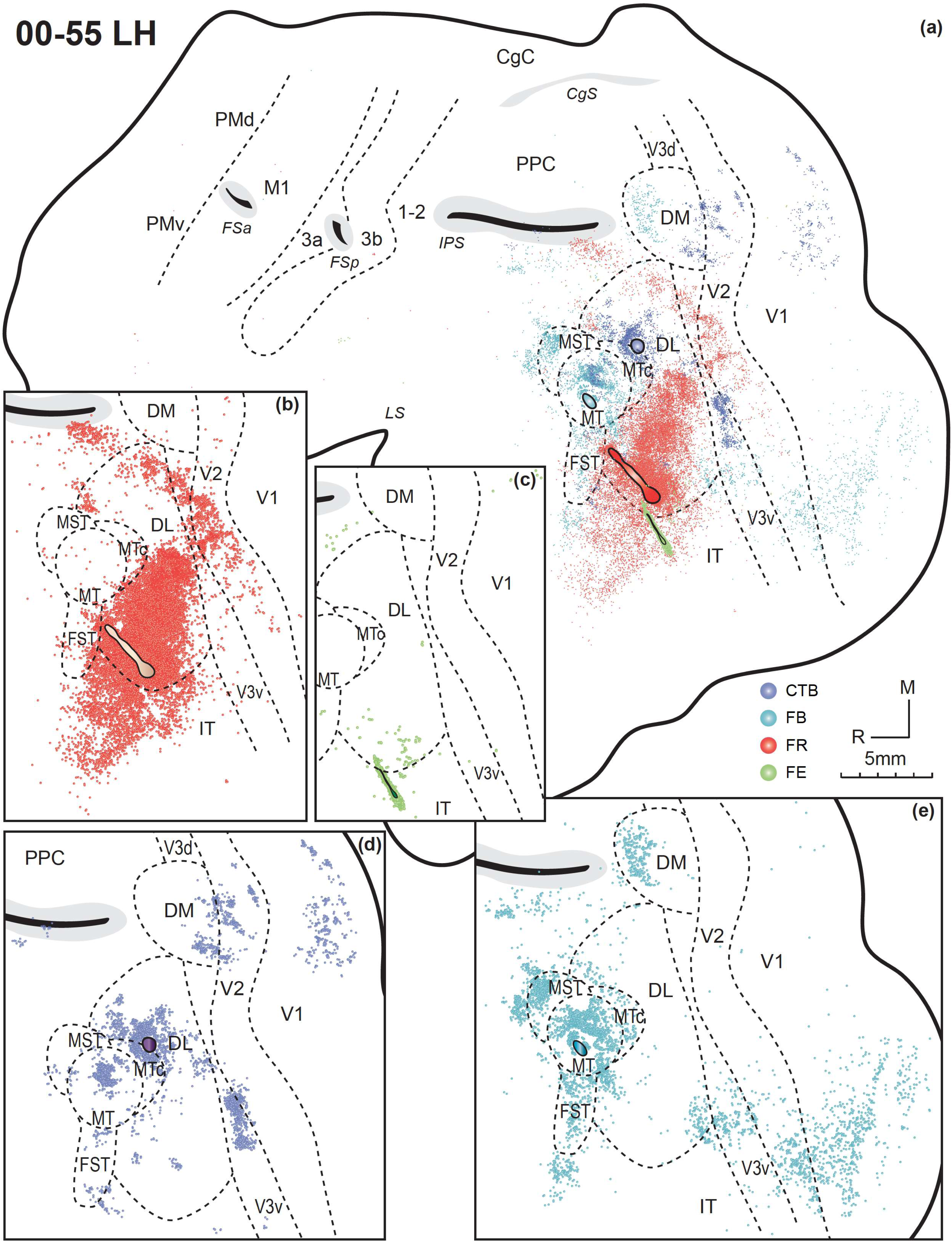
Distributions of labeled neurons after FR, FE, CTB and FB injections in the left hemisphere of galago 00-55, shown altogether in the flattened cortex (a), as well as shown individually by tracers FR (b), FE (c), CTB (d), and FB (e). Other conventions as in Figure 2.

### Case 00-36

Injections of CTB and FR in this case were both placed in visual cortex of the left hemisphere (Figure 6). The FR injection near central DL labeled neurons in ventral and central DL, V2 and V1, as well as ventral V3. Other labeled neurons were scattered in IT and caudal PPC. The CTB injection in IT labeled patches of neurons extending more ventrally in IT, suggesting a series of visual domains in IT.

**Figure 6.**
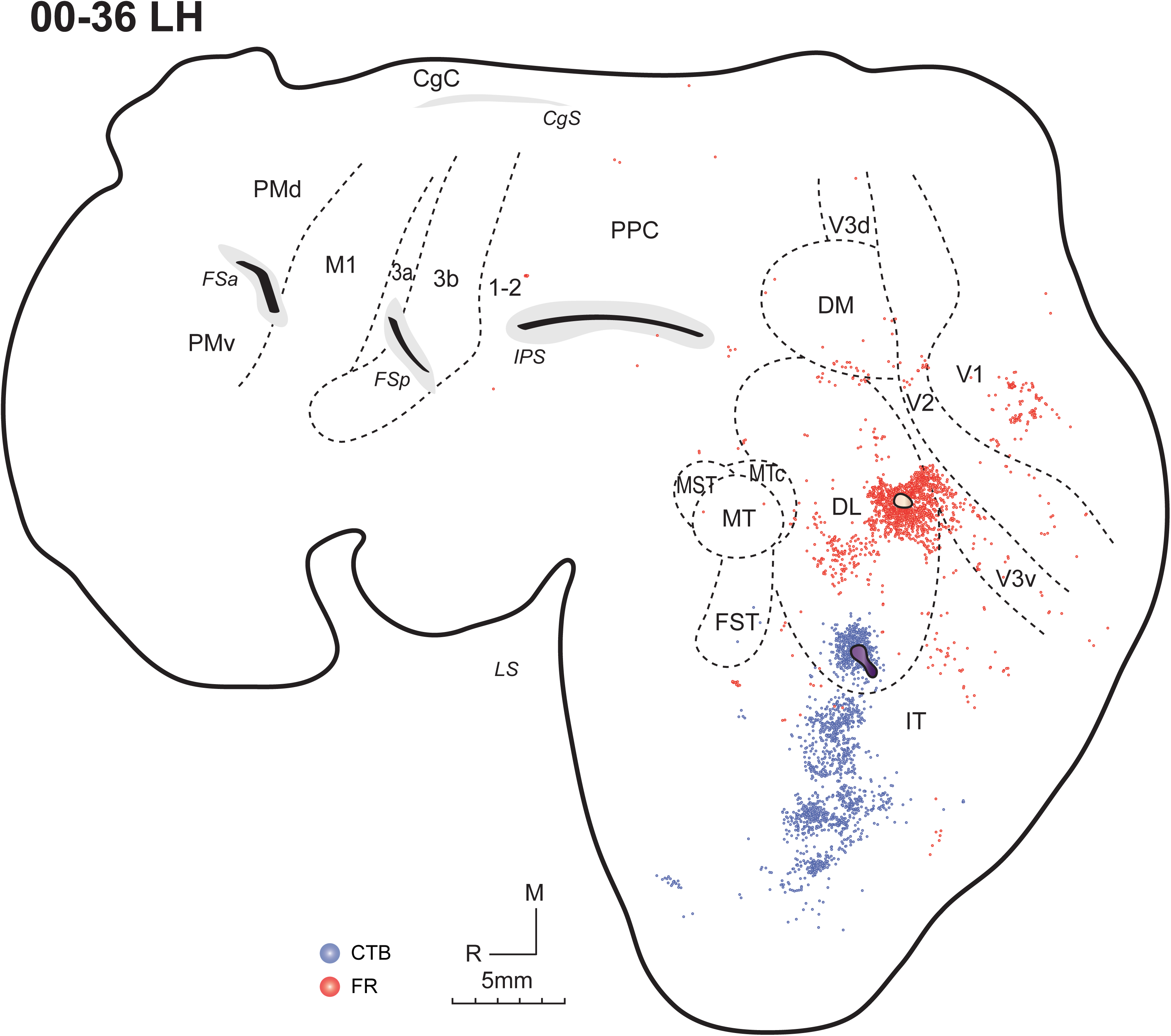
Distributions of labeled neurons after CTB and FR injections in the left hemisphere of galago 00-36, shown altogether in the flattened cortex. Other conventions as in Figure 2.

Overall, the anatomical results from 16 injection sites in visual cortices of five galagos support previous proposals of how the visual cortex is organized in galagos (Figure 7), while suggesting that there are some uncertainties about the extent and possible subdivisions of DL. The proposed DL region has connections with PPC, and MT has connections with DM and PPC. Dorsal DL also receives inputs from IT, which appears to have multiple domains that are visually related.

**Figure 7.**
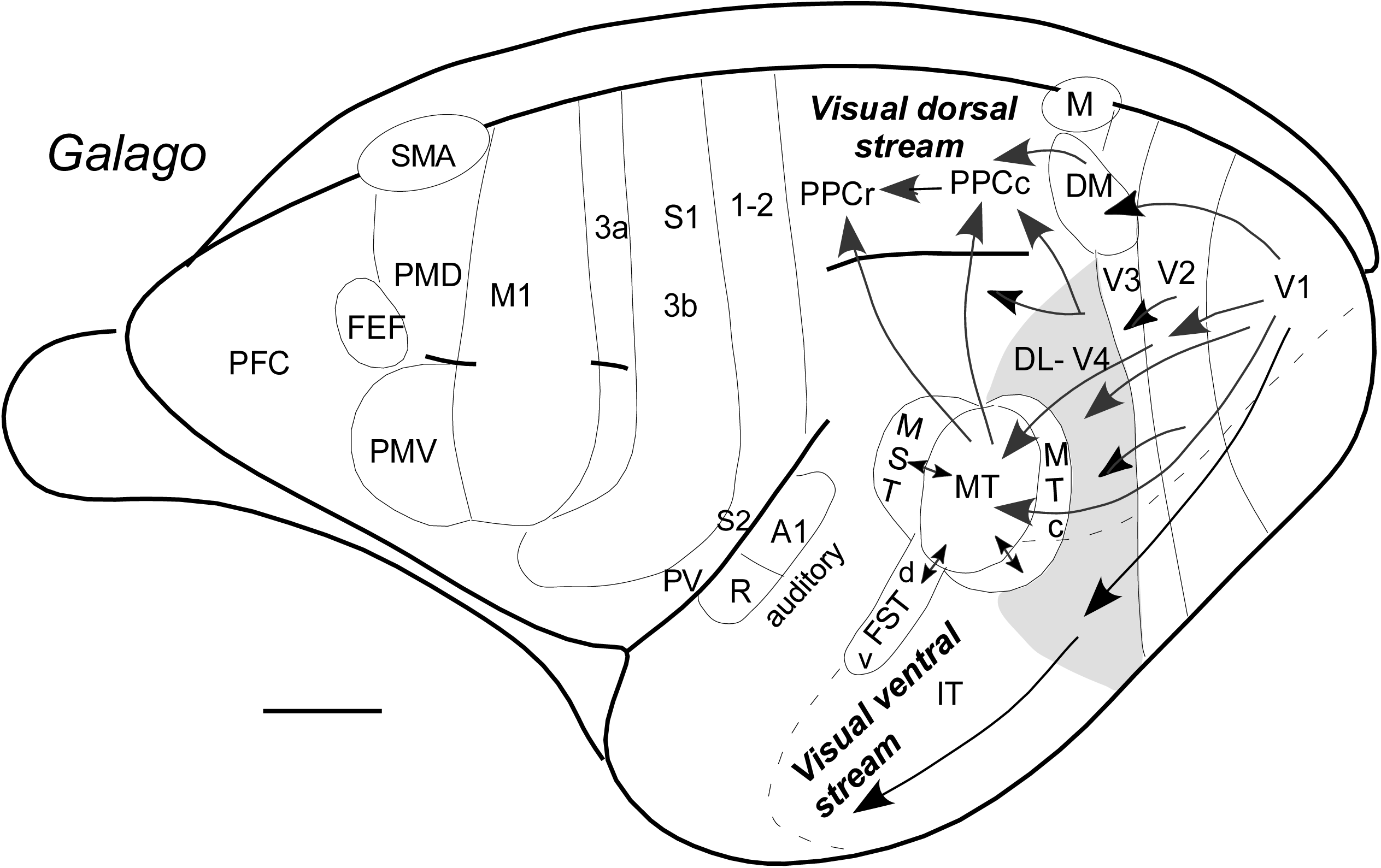
Schematic of proposed subdivisions of cortex in a lateral view of the left cerebral hemisphere of a galago. Visual areas include traditional V1 and V2 areas, a V3 that is disrupted by DM and area M on the medial wall. DL-V4 located between V3 and MT complex is shaded in grey. The MT complex includes MT, MTc, MST and dorsal and ventral division of FST. Present results implicate parts of PPCc and IT cortex in visual processing. Somatosensory areas 3a, 3b-S1, 1-2, PV, and S2 and auditory areas R and A1. Motor areas M1, SMA, PMD and PMV and FEF are shown for reference. LS, lateral sulcus; IPS, intraparietal sulcus; STS, superior temporal sulcus. Scale bar = 5mm.

## Discussion

The present results extend our understanding of visual cortex organization in galagos and reveal the sources of visual inputs to the caudal portion of posterior parietal cortex (PPCc), a component of the dorsal stream of visual processing for visually guided actions (Goodale and Milner, 1992), as well as to inferotemporal cortex (IT), which participates in the ventral stream for object identification (Mishkin and Ungerleider, 1982; Figure 7). The original concept of the two processing streams arose from lesion studies in macaques showing that damage to IT impaired object recognition, whereas lesions of PPC disrupted spatial processing and visuomotor coordination. Subsequent studies in macaques have traced the full extents of these pathways from early visual areas through parietal and temporal cortices to frontal and motor regions.

In the present study, the connectivity patterns of DL in galagos were used to delineate its dorsal and ventral borders and to contribute toward a more unified model of primate visual cortex organization. The connections of DL had not been examined previously in galagos, although their relatively small brains with most visual areas exposed on the cortical surface make them well suited for such investigations. In contrast, DL-V4 connections have been extensively studied in macaques and, to a lesser degree, in New World monkeys including squirrel monkeys, owl monkeys, and marmosets (Wagor et al., 1975; Weller and Kaas, 1987; Steele et al., 1991; Stepniewska et al., 2005; Jeffs et al., 2013). These studies, which employed both tracer injections into the DL–V4 region (Steele et al., 1991; Ungerleider et al., 2008; Stepniewska et al., 2005; Jeffs et al., 2013) and injections into regions projecting to DL-V4 (Wagor et al., 1975; Weller et al., 1984; Weller and Kaas, 1987), generally revealed similar cortical projection patterns across monkeys. DL-V4 consistently connects with V1, V2 and V3, as well as the MT complex, PPC, and IT cortex, with connection density varying by injection site.

Similar patterns were observed here in galagos, though direct comparisons with other primates remain challenging because of differences in proposed cortical parcellations. While there is broad consensus regarding the extent and topography of V1 and V2, the organization of areas immediately rostral to V2 remains controversial. Five different organization schemes have been proposed, and connection patterns may be supportive of more than one model (Angelucci and Rosa, 2015). The main disagreements concern the extent of V3 and the uncertain dorsal and ventral limits of DL-V4. Such inconsistencies complicate cross-species comparisons and highlight the need for additional tract-tracing and visuotopic studies in strepsirrhine species.

Our observations are generally consistent with the widely accepted view that V1, V2, and a modified V3 constitute successive early stages of cortical visual processing in galagos, as in other primates. However, V3d in galagos contains at least one discontinuity filled by an intrusion of DM (Allman and Kaas, 1975; Rosa et al., 1997; Fan et al., 2012). The MT complex, including MT, MTc, MST, FSTd, and FSTv, is well defined in galagos (Kaskan and Kaas, 2007) and in other primates (Kaas and Morel, 1993), and is known to largely contribute to the dorsal stream (Kaas et al., 2022; Figure 7). The DL-V4 region, defined here with approximate dorsal and ventral borders, has incomplete lower visual field representations dorsal to upper visual field representations. DL-V4 may contain several representations, including rostral and caudal subdivisions. Van Essen and Zeki (1978) originally described a “V4 complex” in macaques, suggesting multiple retinotopic representations, and later studies (e.g., Ghose and Ts’o, 1997) indicate that proposed internal functional subdivisions vary across primate taxa.

Our results indicate that DL-V4 in galagos contributes projections to both PPCc and IT cortex. This does not necessarily imply that identical information is transmitted to both dorsal and ventral streams, as little is known about the segregation of functional channels within DL-V4 (Conway and Tsao, 2006). The apparent dorsal border of DL-V4 corresponds to the transition into the upper visual field representation of DM, which likely sets a physiological limit (Stepniewska et al., 2005). The observation that DL-V4 projects to cortex on both sides of the IPS suggests that PPCc in galagos includes at least two or three distinct visual areas, consistent with evidence for multiple visual representations in PPCc of other primates (Felleman and Van Essen, 1991).

The issues raised by these findings relate primarily to the organization of visual cortex in galagos, but they also contribute to a broader comparative framework. Most detailed analyses of visual cortex organization have been conducted in macaques and, to the extent possible, in humans. Other studies in New World monkeys including squirrel monkeys, titi monkey, owl monkeys, and marmosets have clarified additional aspects of cortical organization (Lyon and Kaas, 2002b; Rosa et al., 2005). Galagos, representing a distinct branch of extant strepsirrhines, are of special importance because their visual system and overall cortical organization have been studied more extensively than in any other strepsirrhine species. Galagos are considered as closely resembling the early strepsirrhine euprimates that appeared near the end of the Cretaceous period, around 66 million years ago, possibly preceding the mass extinction that ended the age of dinosaurs (Silcox and Lopez-Torres, 2017; Kaas et al., 2022). Comparative studies of visual system organization in galagos can thus illuminate how extant primates differ and what features were likely present in early primates, offering insights into the evolutionary development of the primate neocortex (for reviews, see Sereno and Allman, 1991; Kaas, 1997; Kaas et al., 2015; Zhu and Vanduftel, 2019).

A particularly controversial issue illustrated in the schematic concerns the nature of area V3 (Figure 7). Early studies in owl monkeys (Allman and Kaas, 1975; 1976) and galagos (Allman et al., 1979) did not identify a complete V3d representing the lower visual quadrant. Instead, regions anterior to V2d appeared to represent the upper visual quadrant, a finding repeatedly confirmed in subsequent studies of galagos (Rosa et al., 1997; Fan et al., 2012) and other primates (Rosa et al., 2005). The present results provide additional support for the view that part of the V3d region represents the upper visual quadrant (Figures 2c, 4b). While some studies have depicted a continuous V3d with a narrow rostral extension of DM, evidence from galagos and other primates supports the conclusion that the upper visual field representation in DM extends to the V2 border (Lyon and Kaas, 2002a; b; c). Thus, the designation of part of the DM territory as V3d needs reconsideration.

Tracer injections in cases 03-40 and 00-78 revealed that DM and adjoining DL both project to PPCc along and posterior to the IPS (Figures 2 and 4). These findings suggest that some injection sites may have extended into DM, but overall, the distribution of labeled neurons within DL–V4 and beyond indicates that the organization of this region cannot be explained by a simple, continuous retinotopic map. Instead, the data suggest a more complex architecture composed of repeating modules representing overlapping portions of visual space. A recent fMRI study in macaques support this view, demonstrating multiple foveal and parafoveal representations within V4 (Wang et al., 2024). If this pattern generalizes, it will require substantial revision of traditional concepts of V4 organization and may explain cross-species differences in the DL-V4 region. Other investigators have subdivided DL-V4 into rostral and caudal regions (Maguire and Baizer, 1984; Steele et al., 1991), extending V3v and adjoining cortex such as VP and VA cortices into portions of the DL-V4 region (Rosa et al., 2005).

Tracer injections into the temporal lobe near or beyond the ventral border of DL–V4 (Figures 3-6) and into dorsal DL-V4 (Figures 2 and 4) further suggest its ventral limits and provide evidence that much of IT cortex participates in higher-order visual processing. The clustering of labeled neurons into three to four distinct zones within IT suggests the presence of multiple, interrelated processing modules in galagos. In other primates, particularly macaques and humans, multiple IT domains specialized for object recognition, including face-selective modules, have been well documented (Arcaro et al., 2020). Overall, these results refine our understanding of visual cortex organization in galagos and support the view that key elements of the dorsal and ventral visual processing streams were already established early in primate evolution.

## Conclusion

The present study contributes to our understanding of visual cortex organization in prosimian primates. Injections of neuroanatomical tracers in the presumptive DL-V4, and adjacent regions of cortex in galagos allowed detailed descriptions of the cortical connections of these regions. Injections of distinguishable tracers were placed in as many as four locations in a single galago, so the connection patterns of several cortical regions could be directly compared in the same animal. The cortical connection patterns of early visual areas support few main conclusions: (1) Patterns of cortical connections of early visual areas V1 and V2 suggest that dorsal V3 has a gap in the representation of the lower visual field that is occupied by the proposed DM area, thus DM gets inputs for upper and lower visual fields from V1 and V2, while V3d handles lower field; (2) Differences in connection patterns of DL and adjacent areas provide evidence for the dorsal and rostroventral limits of the DL-V4 complex; and (3) MT and DM areas provide the major visual inputs to PPC of the dorsal stream with motion and spatial information for guiding actions like reaching and grasping. In summary, the results support the general conclusion that visual cortex organization in galagos is much like that proposed currently in New World and Old World monkeys.

## Acknowledgments

We thank Dr. Christine Collins and Mary Feurtado for help during surgical procedures, and Laura Trice for histological assistance. This work was supported by National Eye Institute grant (EY02686) to J.H.K.

## Data availability

The data that support the findings of this study are available from the corresponding author upon reasonable request.

## Author contributions

Study concept and design: IS, JHK, QW; conducting the experiments: IS; data analysis and illustration: QW; drafting the manuscript: QW, IS, JHK. All authors have full access to the data and take responsibility for the accuracy of the results.

## Abbreviations

Cortical fields 1-2: somatosensory areas 1 and 2
3a: somatosensory area 3a
3b (or S1): primary somatosensory area
A1: primary auditory area
CgC: cingulate cortex
CMA: cingulate motor area
DL: dorsolateral visual area
DM: dorsomedial visual area
FST: fundal superior temporal visual area
IT: inferotemporal cortex
M: medial visual area
M1: primary motor cortex
MST: middle superior temporal visual area
MT: middle temporal visual area
MTc: crescent areas of middle temporal area
PFC: prefrontal cortex
PMC: premotor cortex
PMd: dorsal premotor area
PMv: ventral premotor area
PPC: posterior parietal cortex
PPCc: caudal posterior parietal cortex
PPCr: rostral posterior parietal cortex
S2: secondary somatosensory area
V1: primary visual area
VS: ventral somatosensory area
V2: secondary visual area
V3d: dorsal third visual area
V3v: ventral third visual area
V4: fourth visual area
Sulci CgS: cingulate sulcus
FSa: anterior frontal sulcus
FSp: posterior frontal sulcus
IPS: intraparietal sulcus
LS: lateral sulcus
Anatomical tracers CTB: cholera toxin subunit B
FR: fluororuby
FE: fluoroemerald
FB: fast blue
DY: diamidino yellow

